# Endogenous viral elements reveal associations between a non-retroviral RNA virus and symbiotic dinoflagellate genomes

**DOI:** 10.1101/2022.04.11.487905

**Authors:** Alex J. Veglia, Kalia S.I. Bistolas, Christian R. Voolstra, Benjamin C. C. Hume, Serge Planes, Denis Allemand, Emilie Boissin, Patrick Wincker, Julie Poulain, Clémentine Moulin, Guillaume Bourdin, Guillaume Iwankow, Sarah Romac, Sylvain Agostini, Bernard Banaigs, Emmanuel Boss, Chris Bowler, Colomban de Vargas, Eric Douville, Michel Flores, Didier Forcioli, Paola Furla, Pierre Galand, Eric Gilson, Fabien Lombard, Stéphane Pesant, Stéphanie Reynaud, Shinichi Sunagawa, Olivier Thomas, Romain Troublé, Didier Zoccola, Adrienne M.S. Correa, Rebecca L. Vega Thurber

## Abstract

Endogenous viral elements (EVEs) offer insight into the evolutionary histories and hosts of contemporary viruses. This study leveraged DNA metagenomics and genomics to detect and infer the host of a non-retroviral dinoflagellate-infecting +ssRNA virus (dinoRNAV) common in coral reefs. As part of the Tara Pacific Expedition, this study surveyed 269 newly sequenced cnidarians and their resident symbiotic dinoflagellates (Symbiodiniaceae), associated metabarcodes, and publicly available metagenomes, revealing 178 dinoRNAV EVEs, predominantly among hydrocoral-dinoflagellate metagenomes. Putative associations between Symbiodiniaceae and dinoRNAV EVEs were corroborated by the characterization of dinoRNAV-like sequences in 17 of 18 scaffold-scale and one chromosome-scale dinoflagellate genome assembly, flanked by characteristically cellular sequences and in proximity to retroelements, suggesting potential mechanisms of integration. EVEs were not detected in dinoflagellate-free (aposymbiotic) cnidarian genome assemblies, including stony corals, hydrocorals, jellyfish, or seawater. The pervasive nature of dinoRNAV EVEs within dinoflagellate genomes (especially *Symbiodinium*), as well as their inconsistent within-genome distribution and fragmented nature, suggest ancestral or recurrent integration of this virus with variable conservation. Broadly, these findings illustrate how +ssRNA viruses may obscure their genomes as members of nested symbioses, with implications for host evolution, exaptation, and immunity in the context of reef health and disease.

## Introduction

Endogenous viral elements, or “EVEs,” arise when whole or fragmented viral genomes are incorporated into host cell germlines. Once integrated, EVEs may propagate across successive host generations, potentially becoming fixed in a population through natural selection or drift (Johnson 2015, 2019). Therefore, the presence and content of EVEs can provide clues into the evolutionary relationships among host species and shed light on ancient and modern virus-host interactions (Johnson 2010). To date, most EVEs described in metazoan and plant genomes are retroviral, as this viral group must integrate their genome (as a provirus) into the genome of the host to replicate. Retroviruses thus possess and encode all of the molecular machinery (e.g. reverse transcriptases, integrases) required to integrate autonomously (Stoye 2012). Remarkably, however, sequences from viruses that do not encode reverse transcriptases or exploit integration as a component of an obligate replication strategy – even viruses with no DNA stage – have also recently been detected as EVEs in diverse eukaryotic genomes (Gallot-Lavallée & Blanc 2017, Flynn and Moreau 2019, Horie et al. 2010, Katzourakis & Gifford 2010, Chiba et al. 2011, Chu et al. 2014, Kojima et al. 2021). These non-retroviral RNA EVEs have been reported in hosts ranging from unicellular algae to chiropteran (bat) genomes (Ballinger et al. 2012, Tromas et al. 2014, Palantini et al. 2017, Wang et al. 2014, Jebb et al. 2020, Moniruzzaman et al. 2020, Skirmuntt et al. 2020). Though the mechanisms behind non-retroviral integration continue to be explored, viral sequences may be introduced via nonhomologous recombination and repair, through interactions with host-provisioned integrases and reverse transcriptases supplied on mobile elements (e.g. retrotransposons), or by utilizing co-infecting viruses (Horie et al. 2010, Flynn & Moreau 2019).

Endogenization of any viral sequence (including non-retroviral EVEs) may have positive, neutral or negative effects on a host (Roossinck 2011, Harrison & Brockhurst 2017, Correa et al. 2021). While many EVEs are functionally defective or deleterious and ultimately removed from a population via purifying selection, retained EVEs may remodel the genomic architecture of their hosts or introduce sources of genetic innovation later co-opted for host function (i.e. exaptation; Jern & Coffin 2008, Oliveira et al. 2008). Such ‘domesticated’ EVEs are utilized by hosts as regulatory elements, transcription factors, transposons, and templates for protein coding functions for purposes ranging from developmental pathways to brain function (Feschotte & Gilbert 2012, Frank & Feschotte 2017, Sofuku & Honda 2017, Takahashi et al. 2019). In particular, non-retroviral EVEs potentially serve as antiviral prototypes that help hosts combat infection by exogenous viruses currently circulating in the population (e.g. RNAi; Witfield et al. 2017, Ter Horst 2019, Palantini et al. 2017, Suzuki et al. 2020). If expressed, EVEs may have a significant influence on the health, physiology and/or behavior of their hosts in natural and experimental systems (Parker & Brisson 2019, Suzuki et al. 2020, Wilson et al. 2001).

Investigating the distribution, sequence identity, and function of EVEs can yield insight into virus-host interactions across generations. EVEs catalogue a subset of the viruses that a host lineage has encountered and can link homologous extant viruses to contemporary hosts or known disease states (Holmes 2011, Suzuki et al. 2020). Because integrated elements may accrue mutations at a slower rate than exogenous viral genomes (Aiewsakun and Katzourakis, 2015, Flynn & Moreau 2019), EVEs can fill gaps in virus-host networks and act as synapomorphies, indicating the minimum time that a virus may have interacted with a host. As ‘genomic fossils’, EVEs have helped paleovirologists date the minimum origin of *Circoviridae, Hepadnaviridae, Bornaviridae, Orthornavirae, Lentiviridae*, and *Spumaviridae* infections within metazoans (Feschotte & Gilbert 2012, Johnson 2019). The evolutionary and ecological contexts that EVEs provide for exogenous viruses are particularly informative in understanding the interactions of viruses with multipartite or nested symbiotic systems (Patel et al. 2011, Katzourakis 2013, Aiewsakun & Katzourakis, 2015).

Coral holobionts – the cnidarian animal and its resident microbial assemblage, including dinoflagellates in the family Symbiodiniaceae, bacteria, archaea, fungi, and viruses – are an ecologically and economically valuable, multipartite non-model system (Knowlton & Rohwer, 2003, Matthews et al. 2020). Symbiodiniaceae are key obligate nutritional symbionts of corals and support their hosts in the construction of reef frameworks (LaJeunesse et al. 2018). However, environmental stress can break down coral-Symbiodiniaceae partnerships, resulting in bleaching – the mass loss of Symbiodiniaceae cells (Glynn 1996). Some bleaching signs (paling of a coral colony) are hypothesized to also result from viral lysis of Symbiodiniaceae (van Oppen et al. 2009, Correa et al. 2016, Vega Thurber et al. 2017, Messyasz et al. 2020, Correa et al. 2021, Grupstra et al. 2022), but direct evidence supporting this hypothesis remains limited. Overall, the role of viruses in coral colony health and disease requires further examination.

Non-retroviral +ssRNA dinoRNAV sequences were first reported in stony corals based on five metatranscriptomic sequences and corroborated by Symbiodiniaceae EST libraries (Correa et al. 2013). Subsequent studies indicated that similar +ssRNA viruses are commonly detected in coral RNA viromes and metatranscriptomes, as well as via targeted amplicon assays (Weynberg et al. 2014, Levin et al. 2017, Montalvo-Proano et al. 2017, Grupstra et al. 2022). These viruses exhibit synteny and significant homology to Heterocapsa circularisquama RNA virus (HcRNAV; Levin et al. 2017), the sole recognized representative of the genus *Dinornavirus* and a known pathogen of free-living dinoflagellates (Nagasaki et al. 2005). Both HcRNAV and dinoRNAV sequences detected in coral holobiont tissues contain two ORFs – a Major Capsid Protein (*MCP*) and RNA dependent RNA polymerase (*RdRp*). Furthermore, icosahedral virus-like particle (VLP) arrays resembling HcRNAV (but with 40% smaller individual particle diameters) have been imaged in the Symbiodiniaceae-dense coral gastrodermis tissue and in Symbiodiniaceae themselves (Lawrence et al. 2014). Levin et al. (2017) assembled the 5.2kb genome of a putative dinoRNAV from a poly(A)-selected metatranscriptome generated from cultured *Symbiodinium*. The assembly contained a 5’ dinoflagellate spliced leader (“dinoSL”; Zhang et al. 2013) — a component of >95% of Symbiodiniaceae mRNAs, speculated to illustrate molecular mimicry — and exhibited >1000-fold higher expression in a thermosensitive *Cladocopium C1* population relative to a thermotolerant population of this Symbiodiniaceae strain at ambient temperatures (27°C, Levin et al. 2017, LaJeunesse et al. 2018). Together, the findings from these studies suggest that Symbiodiniaceae are target hosts of reef-associated dinoRNAVs.

This study (1) systematically searched for putative endogenized dinoRNAVs in metagenomes from *in situ* (symbiotic) coral colonies and seawater, as well as in available genomes of Symbiodiniaceae and aposymbiotic (symbiont-free) cnidarians, (2) investigated the evolutionary relationship of putative dinoRNAV EVEs to exogenous reef-associated dinoRNAV sequences, and (3) made preliminary inferences regarding the distribution and possible function of these dinoRNAV EVEs based on their detection, prevalence, and genomic context.

## Methods

### Identification and computational validation of dinoRNAV EVEs leveraging meta’omics

The Tara Pacific Expedition (2016-2018) sampled coral reefs to investigate reef health and ecology using multiple methods, including amplicon sequencing and metagenomics (Planes et al. 2019). In this study, we explored metagenomes generated from hydrocorals (n=60 *Millepora*), stony corals (n=108 *Porites*, n=101 *Pocillopora*) sampled from 11 islands across the South Pacific Ocean during the Tara Pacific Expedition for dinoRNAV EVEs (Figure 1, Supplementary Table 1A, 1B; Pesant et al. 2020). Amplicon libraries of the dinoflagellate Internal Transcribed Spacer 2 (ITS2) gene fragment were sequenced in tandem with the metagenomes, to characterize the dominant Symbiodiniaceae harbored by hydrozoan and stony coral colonies (Hume et al. 2020).

**Figure 1.**
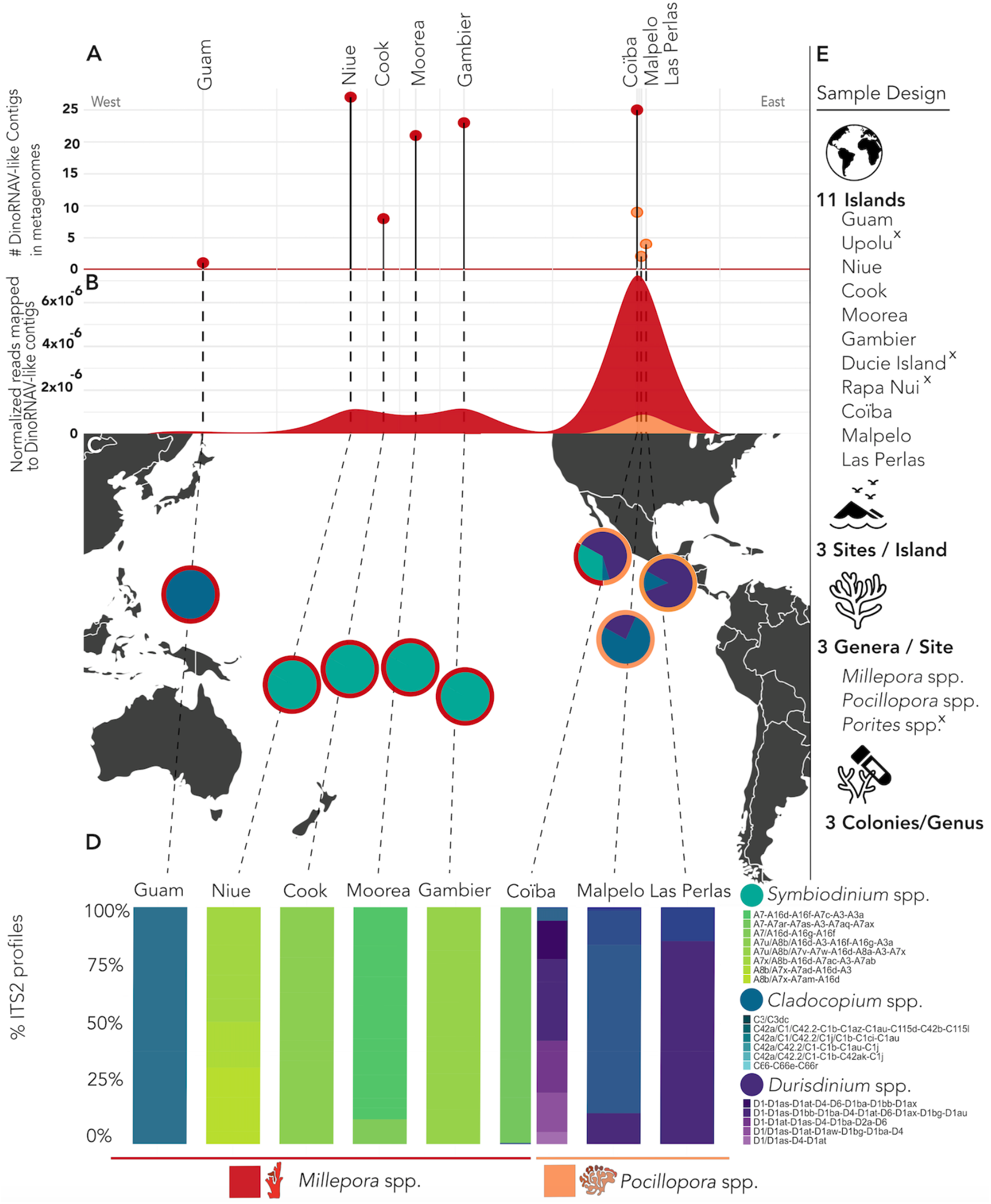
Islands and species (cnidarian and dinoflagellate) correlating with dinoRNAV EVE-like sequence detection among Tara Pacific metagenomes. (A) Count of scaffolds with putative endogenized dinoRNAV-like sequences among Tara Pacific metagenomes, grouped by island and spaced longitudinally by location sampled. (B) Reads mapped to dinoRNAV-like scaffolds within individual Tara Pacific metagenomic libraries, normalized by quality-controlled reads. (C) Sampling sites of Tara Pacific metagenomes explored for endogenized dinoRNAV-like sequences in this study. Internal circles indicate dominant Symbiodiniaceae genera based on ITS2 type profiles, outer ring denotes coral host(s) sampled at each island. (D) Symbiodiniaceae ITS2 type profile metabarcoding as delineated via Symportal (Hume et al, 2019) within island and host. (E) Sample design of Tara Pacific libraries queried for dinoRNAV EVEs. [x] indicates islands or species where no dinoRNAV-like sequences were detected. *[this image is a low resolution placeholder]*

To confirm that these dinoRNAV EVE sequences were affiliated with coral holobionts and reduce the possibility that they are technical artifacts, publicly available metagenome libraries were analyzed. These additional libraries included 120 assembled pelagic water samples presumed to include pelagic dinoflagellate sequences from the Tara Oceans dataset (2009-2013; Pesant et al. 2015) and 30 MiSeq metagenomes from unfractionated samples of the stony coral genus *Acropora*, which were processed and sequenced via a different pipeline (Supplemental Table 1B, Supplemental Figure 1). Publicly accessible transcriptomes from *Symbiodinium microadriaticum* (Supplemental Table 1B) were also queried to determine if dinoRNAV-like sequences were present in poly(A)-selected dinoflagellate transcriptomes and resembled EVEs in terms of proximal gene composition and presence of a characteristic pre-mRNA spliced leader (SL) sequence (as in Levin et al, 2017). Details regarding the collection of samples, generation of metagenomes and associated Symbiodiniaceae amplicon libraries, and associated bioinformatic analyses are provided in Supplementary Figure 1).

Metagenomic and transcriptomic scaffolds were annotated against a curated database of dinoRNAV-like sequences (Supplemental Table 2) via BLASTx (Altschul et al. 1990, Supplementary Figure 1). Alignments to the custom database with a bit score <50 and percent shared amino acid identity <30% were excluded from further analysis. A length penalty was not imposed during this step due to the limited length of assembled scaffolds (average N50=3341±127 nt across all queried libraries). Open reading frames (ORFs) from selected scaffolds were called via Prodigal (v.2.6.3; Hyatt et al. 2010) and annotated against the NCBI-nr database (DIAMOND v.2.0.6; Buchfink et al. 2015) to confirm homology to dinoRNAVs and identify adjacent dinoflagellate sequences. In the absence of complete ORFs (potentially due to the limited size of scaffolds, partial integrations, etc.), homology was confirmed through comparison of the initial alignments to the curated database and 300nt of upstream/downstream flanking sequences (bedtools v.2.30.0; Quinlan et al, 2010) against the NCBI-nr database. Non-normalized quality-controlled reads were mapped via bbmap (v.38.84; Bushnell et al. 2017), and putative EVEs were assessed for uniform read coverage across scaffolds, reducing the likelihood of chimeric assembly. RNA secondary structure was predicted via mfold (v.3.5; Zuker et al 2003).

### dinoRNAV EVEs in dinoflagellate and aposymbiotic cnidarian genomes

Publicly available dinoflagellate and aposymbiotic (dinoflagellate-free) cnidarian genome assemblies were queried to resolve the putative host(s) of dinoRNAVs, to assess homology among detected dinoRNAVs within coral holobionts, and to compare genes proximal to dinoRNAV EVEs in different host species/strains. A chromosome-scale dinoflagellate genome assembly generated from a *Symbiodinium microadriaticum* culture (Accession: GSE152150, Nand et al. 2021), and scaffold-scale genome assemblies were examined for dinoRNAV EVEs (Supplemental Table 1B). Scaffold-scale genome assemblies were from the closely related families Symbiodiniaceae and Suessiaceae, and included representatives from the genera *Symbiodinium* (n=9), *Breviolum* (n=1), *Cladocopium* (n=3), *Durusdinium* (n=1), *Fugacium* (n=2), and *Polarella* (n=2), as well as 25 aposymbiotic cnidarian genome assemblies, including the stony coral genera *Acropora* (n=13), *Astreopora* (n=1), *Galaxea* (n=1), *Montastraea* (n=1), *Montipora* (n=3), *Orbicella* (n=1), *Pocillopora* (n=2), *Porites* (n=1), and *Stylophora* (n=1), and the jellyfish *Clytia* (n=1; Figure 2, Supplemental Table 1B). Genome completeness and quality were assessed via BUSCO (v3; Simão et al. 2015) with the Eukaryota dataset and QUAST (v5.0.2; Gurevich et al. 2013), respectively. Scaffolds/chromosomes containing putative dinoRNAV EVEs were identified by aligning sequences to the protein version of the Reference Viral DataBase (RVDB v.19; Bigot et al. 2019) using DIAMOND BLASTx (v0.9.30; Buchfink et al. 2015). The same exclusion criteria were maintained for alignments of metagenomic scaffolds, also omitting alignments <100 amino acids. Regions of dinoflagellate genomes exhibiting similarity to the *MCP* or *RdRp* of reef-associated dinoRNAV reference genomes (Levin et al. 2017) or other closely related +ssRNA viruses (Supplemental Table 2) were extracted and re-aligned to the NCBI-nr database to further confirm viral homology. Putative whole dinoRNAV-like genomes within scaffolds were identified based on the presence of *MCP* and *RdRp-like* sequences on the same scaffold no further than 1.5 Kbp apart (Table 1; Supplemental Figure 2). IRESPred (Kolekar et al., 2016) was utilized to identify internal ribosomal entry sites (IRES) on putative dinoRNAV EVE with whole sequence integrations.

**Figure 2.**
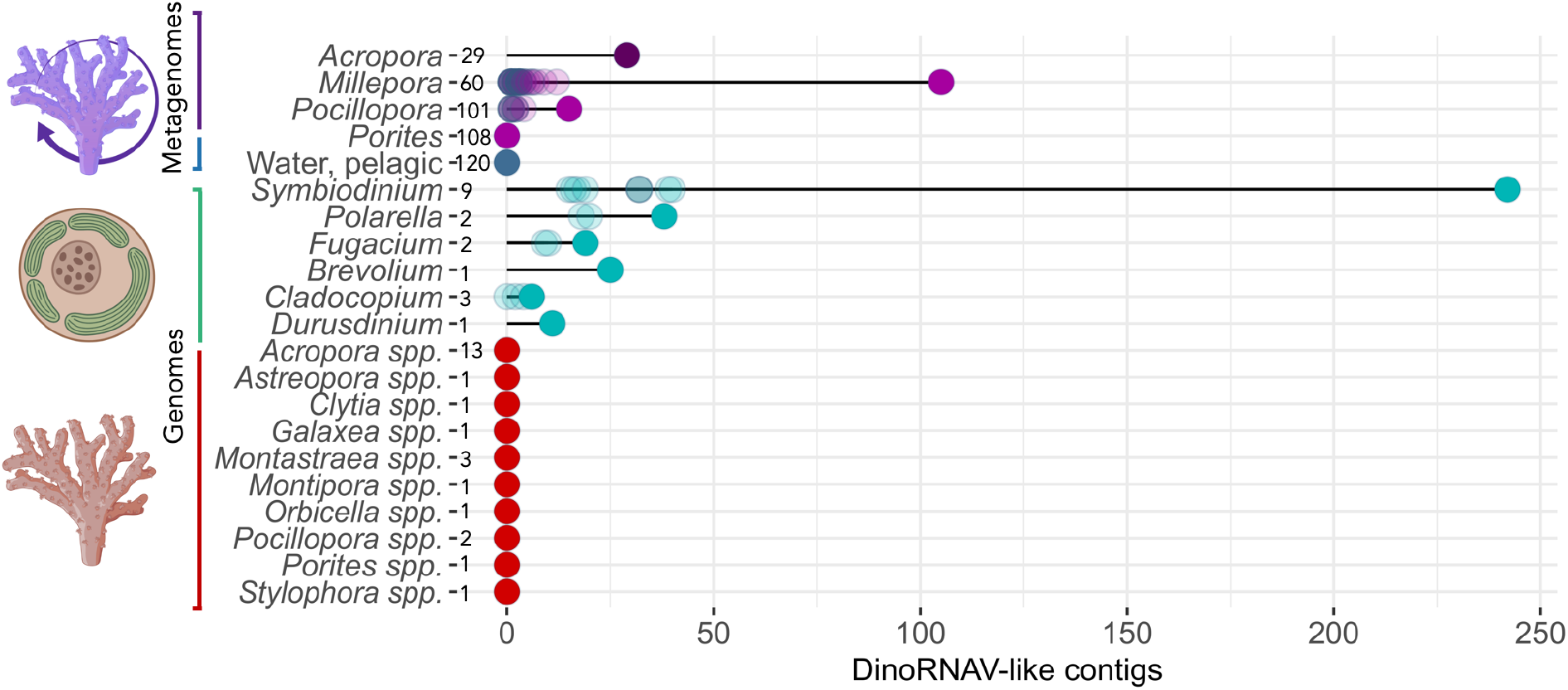
Total quantity of putative endogenized dinoRNAV EVEs identified, broadly organized by sample source (metagenome or genome), and number of libraries or assemblies queried (numbers follow a dash to the right of source name). Opaque circles denote the sum total of dinoRNAV EVE-like sequences identified from each source, while transparent circles denote independent counts per library queried.

ORFs were predicted and annotated from dinoRNAV EVE-containing scaffolds and all dinoflagellate chromosomes using Prodigal (Hyatt et al. 2010) and MAKER2 annotation pipeline (Holt and Yandell 2011) with the AUGUSTUS gene prediction software (Stanke et al. 2006). Translated ORFs were then aligned to a hybrid database containing the UniProt/Swiss-Prot database and protein version of RVDB (v.19; DIAMOND-BLASTp). ORFs on putative dinoRNAV EVE-containing scaffolds and chromosomes were further annotated using InterProScan (v5.48-83.0, Pfam, PANTHER) to identify sequences proximal to putative dinoRNAV integrations.

### Phylogenetic analysis of dinoRNA VEVEs

Amino acid-based phylogenetic trees were generated with dinoRNAV EVE ORFs (*MCP* and *RdRp*) from scaffold-scale genomic assemblies, metagenomes, transcriptomes, and sequences from exogenous and closely related +ssRNA reference viruses (Supplemental Table 1A,B, Supplementary Table 2). Sequences were aligned using the best fit algorithm determined by MAFFT (v7.464; Katoh and Standley 2013) and reviewed and trimmed manually in MEGA (v7; Kumar et al. 2016). Maximum-likelihood trees were generated with IQTREE2 (Minh et al. 2020) using the model determined by ModelFinder (Kaylaanamoorthy et al. 2017) and 50,000 parametric bootstraps (Hoang et al. 2018) with nearest neighbor interchange optimization.

## Results and Discussion

### Evidence of Endogenized dinoRNAVs in Coral Holobiont Metagenomes

Putative dinoRNAV EVEs were detected in metagenomes generated from 42 cnidarian holobionts out of 269 sampled across the South Pacific Ocean. The majority of endogenized dinoRNAVs were identified in hydrocoral metagenomes (*Millepora* spp.) which predominantly harbored *Symbiodinium* dinoflagellates (n=105; 70.5%), but EVE-like sequences were also observed in scleractinian coral metagenomes (*Pocillopora* spp.) which predominantly harbored *Cladocopium* and *Durusdinium* dinoflagellates (n=15; 29.5%; Figure 1B,C). No dinoRNAV-like sequences were detected among *Porites* spp. metagenomes (Figure 1, Figure 2). Hydrocoral metagenomes were sequenced at equivalent depths as scleractinian corals and had comparable levels of annotation (Supplementary Table 3); thus, higher dinoRNAV EVE prevalence in hydrocoral libraries was likely not a result of methodological bias. Of the 11 evaluated South Pacific islands, dinoRNAV EVEs were identified in samples from eight (Guam, Gambier, Moorea, Cook, Niue, Malpelo, Coïba, and Las Perlas), spanning 18 unique sites (Figure 1C). There was a distinct longitudinal trend among *Pocillopora* spp. metagenomes; putative dinoRNAV EVEs were only identified in this coral genus on the Central American coast (CAMR, Coastal Pacific Longhurst Province).

Importantly, endogenized dinoRNAV open reading frames (ORFs) appeared to be immediately adjacent to ORFs identified as dinoflagellate (typically Symbiodiniaceae) genes— they were not proximal to coral genes or those of other cellular organisms abundant in these metagenomes (Supplemental Table 4). We examined the Symbiodiniaceae ITS2 profiles (Hume et al. 2020) associated with each metagenome and found that putative dinoRNAV EVEs were primarily associated with *Symbiodinium, Cladocopium*, and *Durusdinium*, which exhibited variation on both host and regional scales (Figure 1D). DinoRNAV EVEs were more common in *Symbiodinium*-dominated cnidarians (*F_2,1044_*=*25.8*, p<0.0001, nested ANOVA; Supplemental Figure 3) relative to cnidarians hosting other Symbiodiniaceae genera, regardless of host. This suggested that dinoRNAV integration may be particularly recurrent or conserved within the genus *Symbiodinium* (Figure 1).

To determine if these putative viral integrations were specific to cnidarian holobiont metagenomes and ensure that they were not artifacts of shared sample processing and sequencing procedures of the Tara Pacific pipeline, we also analyzed seawater metagenomes and publicly available metagenomes from the stony coral-dinoflagellate holobiont, *Acropora* spp. (Supplemental Table 1B). Examination of 120 Tara Oceans pelagic seawater metagenomes (Pesant et al. 2015) yielded no sequences sharing homology to dinoRNAVs. The concentration of Symbiodiniaceae cells within cnidarian tissues is significantly higher than that of the surrounding seawater (Littman et al. 2008, Schuefen et al. 2017, Fujise et al. 2021, Grupstra et al. 2021). Thus, lack of detection of dinoRNAV-like sequences from seawater metagenomes is likely due to reduced genomic signal of Symbiodiniaceae in the water column. Analysis of the 30 non-Tara *Acropora* holobiont metagenomes identified 29 more putative dinoRNAV EVEs (Figure 2). These dinoRNAV EVEs were again neighboring dinoflagellate ORFs. While the Caribbean *Acropora* metagenomes analyzed contained too few reads to resolve the dominant Symbiodiniaceae present, earlier studies of the same coral colonies identified *Symbiodinium* spp. as the primary symbiont present (Muller et al. 2018).

The identification of endogenized dinoRNAV-like sequences in cnidarian holobiont metagenomes, combined with the proximity of dinoRNAV-like ORFs to dinoflagellate-like sequences across metagenomes harboring diverse dinoflagellate consortia, collectively indicate that dinoRNAV EVEs are widespread among Symbiodiniaceae genera (Figure 2 cyan dots).

### Endogenized DinoRNAVs Detected in Symbiodiniaceae Genomes

To further test the hypothesis that dinoRNAVs on reefs infect dinoflagellate symbionts and not cnidarians, we examined 18 scaffold-scale genome assemblies representing the dinoflagellate families Symbiodiniaceae and Suessiaceae as well as 25 cnidarian genomes spanning 10 genera (Supplemental Table 1B; Figure 2; Table 1). Alignments revealed no evidence of endogenized dinoRNAVs in any of the 151,782 aposymbiotic (dinoflagellate-free) cnidarian scaffolds. In contrast, the same approach uncovered 351 (of 593,433) dinoflagellate scaffolds with evidence of endogenized dinoRNAVs (Figure 2; Table 1). The identified 351 dinoRNAV EVE-containing scaffolds were observed across 17 of the 18 dinoflagellate genome assemblies (Table 1). DinoRNAV EVEs were also observed in two assemblies from the free-living dinoflagellate genus, *Polarella* (family Suessiaceae), which is closely related to the family Symbiodiniaceae, and served as an outgroup in this study (Janouškovec et al. 2017; Stephens et al. 2020). Interestingly, assemblies belonging to *Symbiodinium*, the most ancestral Symbiodiniaceae genus (LaJeunesse et al. 2018), contained a higher number of scaffolds with putative dinoRNAV EVEs (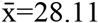, stdev=10.7) relative to assemblies of other Symbiodiniaceae genera (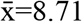, stdev=11; Figure 2 cyan dots; Table 1). This result may clarify why observations of dinoRNAV-like ORFs were more common in metagenomes dominated by *Symbiodinium* (Figure 1D). The dinoflagellate genome assembly with no detected dinoRNAV EVEs belonged to a relatively incomplete assembly of *Cladocopium* C15, which had the second lowest N50 and lowest BUSCO completeness score of all genomes examined (completeness 11.6%, relative to the average 24.54%; Table 1, Supplementary Table 5). The lower coverage/completeness of the *Cladocopium* C15 assembly indicates a reduced window into this genome. It is therefore possible that when a more complete assembly is generated, dinoRNAV EVE-like sequences will be detectable from this dinoflagellate. However, a linear model suggested that there was no relationship between dinoRNAV EVE detection and assembly statistics (i.e. number of scaffolds, N50, completeness). Furthermore, since we were unable to detect dinoRNAV EVEs in *Porites* metagenomes – a coral species primarily harboring *Cladocopium* C15 symbionts – we hypothesize that dinoRNAV endogenization is less common in this lineage of Symbiodiniaceae.

**Table 1.**
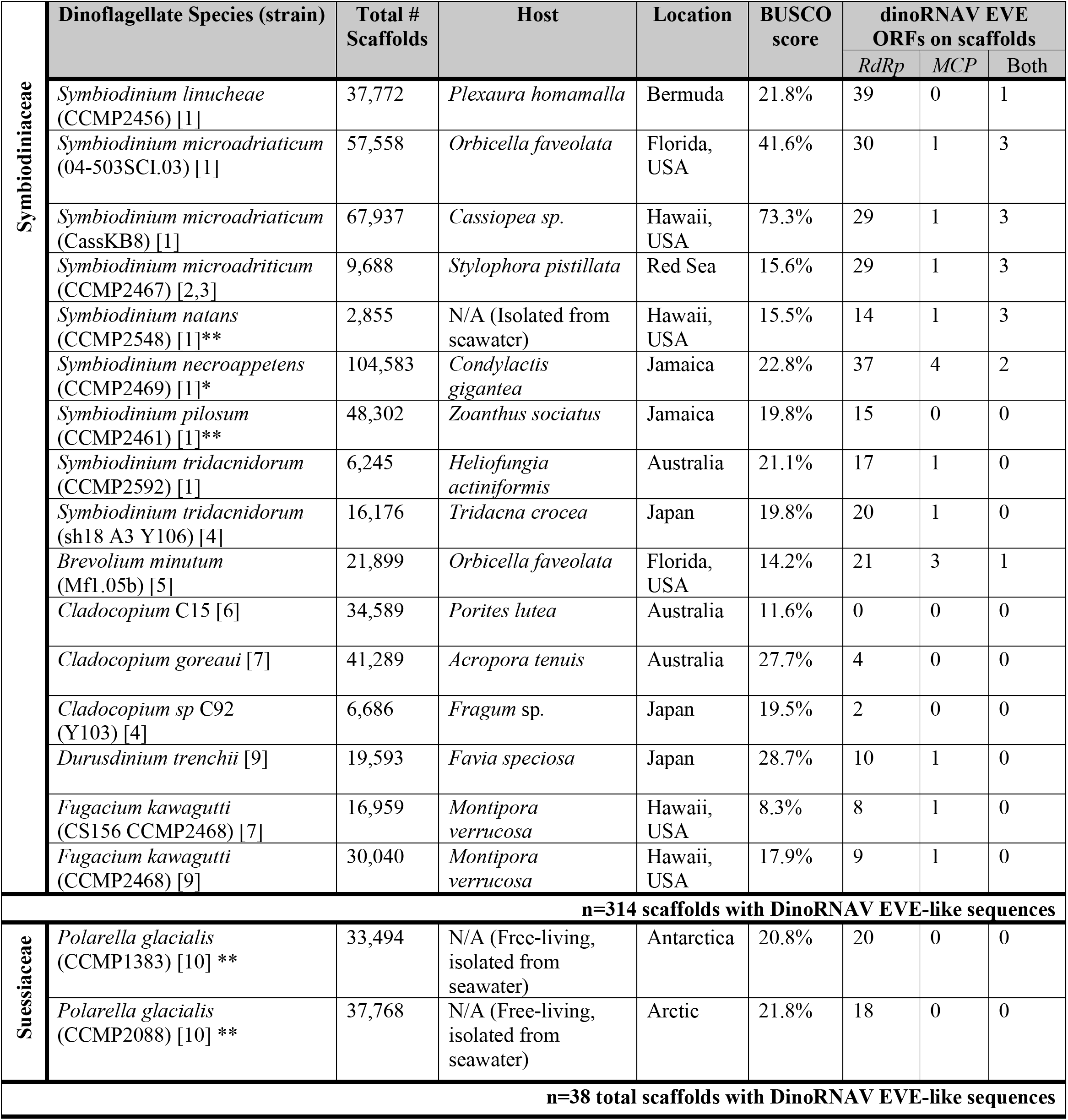
DinoRNAV EVE-like detections from representative Symbiodiniaceae and Suessiaceae dinoflagellate scaffold-level genome assemblies, as well as the host species and location of isolation for each dinoflagellate. Assembly coverage and completeness are measured via BUSCO score (% completeness, or %C; Simão et al, 2015). * Indicates species with documented opportunistic life history; ** Indicates species with documented free-living life history per principal species description. Total counts of dinoflagellate scaffolds in genomes queried with individual endogenized dinoRNAV ORFs (*RdRp, MCP*) or both ORFs nearby one another are provided. *RdRp* = RNA-dependent RNA polymerase; *MCP* = major capsid protein. [1] Gonzalez-Pech et al. 2021, [2] Aranda et al. 2016, [3] Gonzalez-Pech et al. 2019, [4] Shoguchi et al. 2018, [5] Shoguchi et al. 2013, [6] Robbins et al. 2019, [7] Liu et al. 2018, [8] Shoguchi et al. 2020, [9] Lin et al 2015, [10]Stephens et al. 2020. Further genome citations (including accession numbers) and BUSCO completion metrics can be found in Supplementary Table 5.

### Incomplete ORFs and Possible Duplications Indicate Endogenization of DinoRNAVs

The repeated observation of putative dinoRNAV EVEs in dinoflagellate scaffolds and contigs from metagenomes and genomes suggests these sequences are either (1) conserved sequence artifacts of Symbiodiniaceae-dinoRNAV interactions, and/or (2) evidence of highly prevalent dinoflagellate viruses, commonly integrated and propagated via their single-celled hosts. If the observed dinoRNAV-like sequences represent active infections capable of generating virions during egress, we would, at minimum, expect essential ORFs associated with replication (RNA-dependent RNA polymerase, *RdRp*) and virion structure (Major Capsid Protein, *MCP*) to be endogenized on the same scaffold. We would additionally expect to observe overall conservation of ORF length/composition (with a lack of internal stop codons or significant deletions) when aligning the dinoRNAV-like sequences detected here with known exogenous dinoRNAV sequences.

However, both DIAMOND and gene prediction analyses generally depicted dinoRNAV-like ORFs in isolation on separate scaffolds. While 28 MCP and 73 RdRp dinoRNAV ORFs were annotated, both ORFs were present on a Symbiodiniaceae scaffold – potentially representing whole dinoRNAV genome integrations – in only 14 instances. Thirteen of these 14 were from *Symbiodinium* genomes, whereas one scaffold was from *Breviolum minutum*, a member of the second most ancestral dinoflagellate genus (Table 1; LaJeunesse et al. 2018). To assess the conservation of putative dinoRNAV EVE sequence length/composition, we aligned the genomic and single ORF EVEs to reference exogenous dinoRNAV sequences. The reference genome for reef-associated dinoRNAVs is ~5 Kbp long and contains a 1,071 bp noncoding region between ORFs, with a 124-nucleotide internal ribosomal binding site (Levin et al. 2017). In this study, for 13 of the scaffolds in which dinoRNAV ORFs were detected, the putative noncoding region between the *MCP* and *RdRp* EVEs ranged from ~200-800 bp (except for a scaffold belonging to *S. linucheae* CCMP2456, which contained a ~79 kbp noncoding region, and was excluded in further alignments). No internal ribosomal binding sites were detected within the putative dinoRNAV EVEs identified in dinoflagellate genomes. A nucleotide-based alignment to Levin et al.’s (2017) reference dinoRNAV genome indicated that the putative dinoRNAV EVEs presented here contained substantial insertions and/or deletions (Supplemental Figure 2). Translated exogenous dinoRNAV *MCP* ORFs are reported to be ~358 aa in length (Levin et al. 2017; Figure 3 top sequences), but dinoRNAV-like *MCP* sequences recovered in this study ranged from 116-605aa in length. Furthermore, comparisons of these endogenous *MCPs* to exogenous reference sequences revealed internal stop codons and overall low similarity (Figure 3), instead sharing high conservation in structural amino acid motifs. Amino acid-based alignment of endogenous dinoRNAV MCPs to metatranscriptome- and amplicon-generated exogenous reference sequences (Levin et al. 2017, Montalvo-Proaño et al. 2017) revealed that indels and regions of low similarity were observed between three conserved regions across both endogenous and exogenous MCP sequences (red boxes in Figure 3).

**Figure 3.**
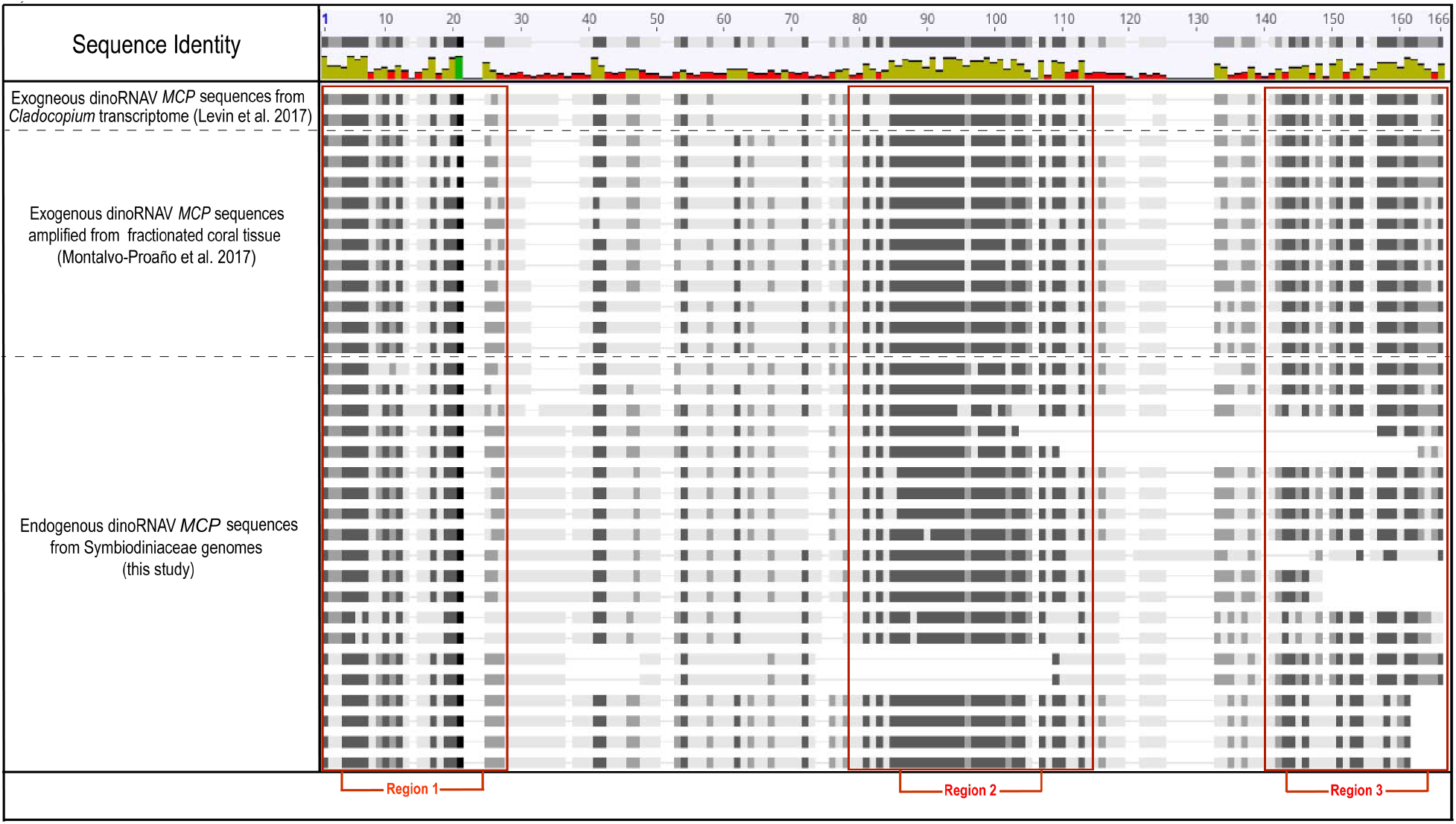
Amino acid alignment including major capsid protein (*MCP*) reference sequences from exogenous dinoRNAV-like +ssRNA viruses, as well as putatively endogenous dinoRNAV *MCP* sequences. Exogenous reference sequences include: 1. Symbiodiniaceae +ssRNA virus *MCP* ORFs recovered from a *Cladocopium* sp. transcriptome (Levin et al, 2017), and 2. dinoRNAV *MCP* amplicons from fractionated coral tissue (Montalvo-Proaño et al. 2017). Conserved regions observed in exogenous and putatively endogenous sequences are labeled as Regions 1-3.

Interestingly, multiple whole dinoRNAV integrations were sometimes observed in a single dinoflagellate genome. For example, genome assemblies of four different *S. microadriacticum* strains contained two or three whole dinoRNAV EVEs each (Table 1; Figure 3).

Pairwise alignments measuring shared nucleotide identity of whole dinoRNAV EVEs across Symbiodiniaceae scaffolds revealed that the *S. microadriaticum* genomes and the *S. necroappetens* genome share two whole genome dinoRNAV EVEs (provisionally dinoRNAV-A and dinoRNAV-B; Supplemental Figure 2; Clustal-Omega; Sievers et al. 2011). *S. microadriaticum* dinoRNAV-B was identical in all strains and shared 97% identity with the *S. necroappetens* dinoRNAV-B, yet proximal genes varied (Supplemental Table 6). The inconsistent composition and fragmented nature of both the genomic and single ORF dinoRNAV EVEs reported here supports the hypothesis that these sequences are not capable of generating replicative virions and are best interpreted as multiple integrations of dinoRNAVs into a host genome.

### A Potential Mechanism for dinoRNAV Endogenization: Host-Provisioned Retroelements

To assess if general genomic “neighborhoods” are conserved across dinoRNAV integrations (e.g. site location and synteny) and to better understand the genes proximal to EVEs on Symbiodiniaceae genomes, a chromosome-scale *Symbiodinium microadriaticum* genome assembly was evaluated (Figure 4). The highest quality dinoflagellate genome assembly currently available revealed dinoRNAV-like ORFs on 18 of 94 chromosomes, with at least one *RdRp* on each, and some with multiple (two with n=2 *RdRps*, three with n=3 *RdRps*). On three of the chromosomes (# 30, 35, and 74), there were predicted ORFs annotated as dinoRNAV *MCPs* in close proximity to a *RdRp* ORF (separated by noncoding regions 319-656nt), indicative of a potential full-length dinoRNAV genome integration. These results corroborate detections of multiple genomic dinoRNAV EVEs in scaffold-scale assemblies of *Symbiodinium microadriticum* genomes (Supplemental Figure 2). The higher-resolution *S. microadriaticum* chromosome-level assembly facilitated the identification of an additional dinoRNAV genomic EVE (n=4 for chromosome-level vs. n=3 for scaffold-level, Supplemental Figure 2), two of which were identified on Chromosome 74 and were separated by 2,501 nucleotides. Of note, Nand et al. (2021) reported a decreasing abundance and expression of genes towards the center of chromosomes (past ~2Mpb of a telomere), where there was an increase in repetitive elements; this is where 26 of 29 putative dinoRNAV EVEs were identified in the chromosome-level assembly. Furthermore, ORFs neighboring integrations often varied widely both in proximity and predicted function: from collagen and RNA binding protein to reverse transcriptase and non-LTR retrotransposable elements, these ORFs potentially contributed to the integration of the putative dinoRNAV EVEs.

Host-provisioned retroelements are a proposed mechanism of non-retroviral RNA virus endogenization (Horie et al, 2010, Flynn & Moreau 2019). We examined the annotated ORFs surrounding dinoRNAV-like sequences on the 18 EVE-containing *S. microadriaticum* chromosomes to gain insight into the integration of dinoRNAVs. *Symbiodinium* contain numerous long interspersed nuclear elements (LINEs) relative to other Symbiodiniaceae genera, with LINES comprising 74.10-171.31 Mbp of *Symbiodinium* genomes, relative to an average of 7.48 Mbp of the genomes of in other genera (González-Pech et al. 2021, Nand et al. 2021, Mita and Boeke 2016). This group of non-long terminal repeat (non-LTR) eukaryotic retrotransposons contains reverse transcriptases and is conserved, implying that they are active and may facilitate non-retroviral endogenization of dinoRNAVs within Symbiodiniaceae. Proximal to putative dinoRNAV *MCP* and *RdRp* ORFs on *S. microadriaticum* chromosomes, ~40% of annotated ORFs (35 of 88 annotated proteins) were similar to non-LTR retrotransposable elements seen in other eukaryotic genomes (Figure 4, Supplemental table 6), sometimes <300bp 5’ upstream. We also annotated ORFs similar to hypothetical virus proteins, suggesting that this mechanism may facilitate integration of sequences beyond dinoRNAVs. These observations of non-LTR retrotransposable elements, sometimes in very close proximity to dinoRNAV EVEs, supports the hypothesis of cis-acting host-driven integration and may explains the increased abundance of dinoRNAV EVEs, particularly in *Symbiodinium* genomes.

**Figure 4.**
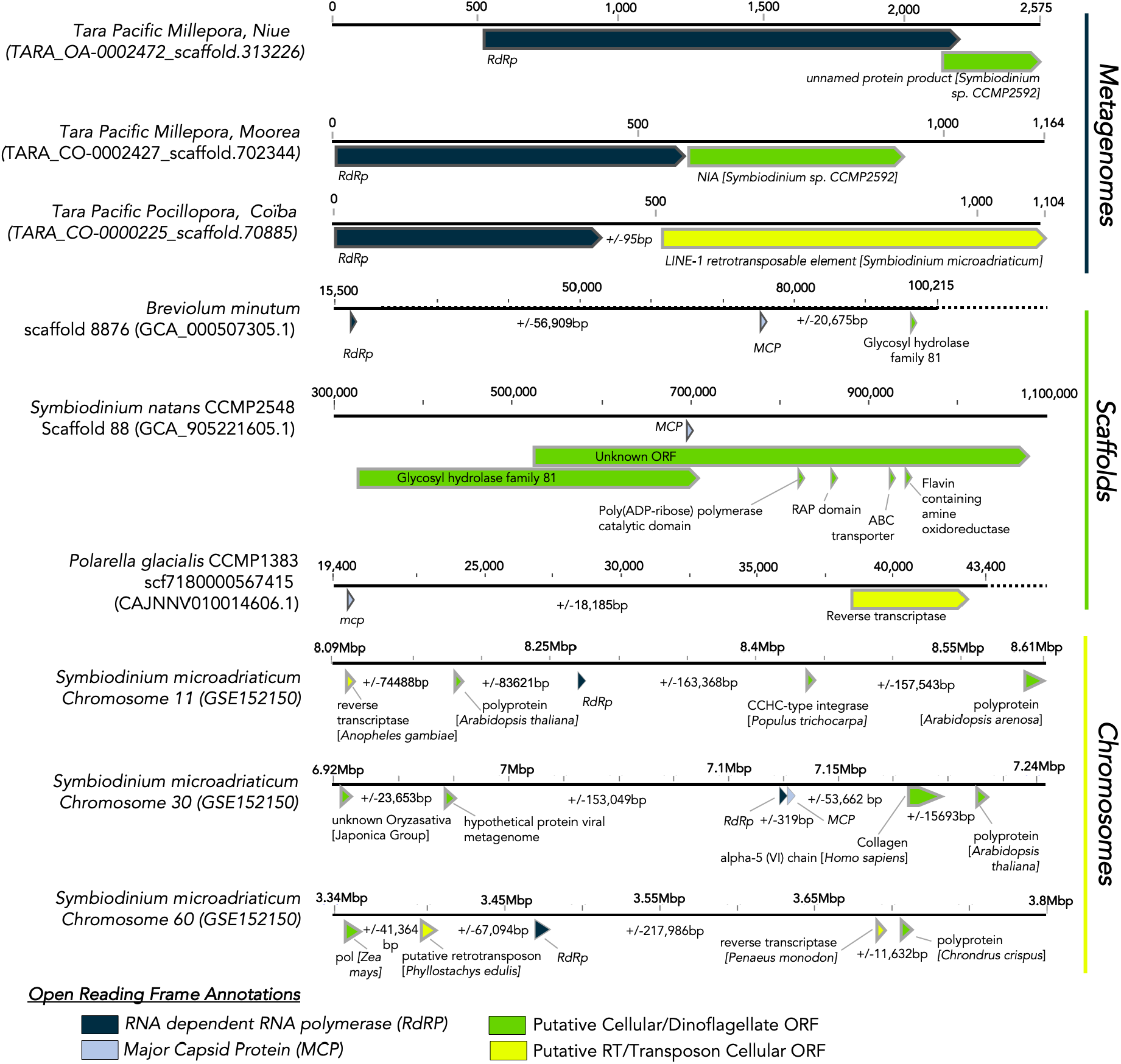
Representative scaffolds and chromosome fragments containing putative dinoRNAV EVEs (*MCP*, light blue; *RdRp*, navy blue ORFs with complete description in supplemental table 6). Open reading frame (ORF) color broadly indicates cellular versus putative +ssRNA viral homology; yellow ORFs may be exploited mechanisms for viral integration. (+/−) base pair values represent sequence lengths between ORFs.

### DinoRNAV EVEs Show Homology to Extant Exogenous Viruses

Endogenized viruses can lend insight into the dynamics between cellular hosts and extant, free virions. Because many EVEs evolve at the rate of the host genome, rather than at the much faster rate of exogenous +ssRNA viral genomes, EVEs can serve as a snapshot of viral ancestry (Holmes et al. 2010). We compared translated putative dinoRNAV EVEs from this study to putative exogenous dinoRNAVs and extant *Dinornavirus* taxa to better understand if the former were highly conserved sequences (with potential utility to the host) and/or recently integrated EVEs. We found that amino acid translations of endogenous dinoRNAV *MCP* sequences contained conserved motifs observed in the exogenous *MCP* sequences (e.g. regions 1-3 in Figure 3), yet the associated phylogeny was highly polyphyletic along inferred ancestral nodes (Figure 5A, B). Several putative dinoRNAV EVEs shared similarity to extant *MCPs* identified from unfractionated stony coral holobionts via amplicon sequencing (Montalvo-Proaño et al. 2017); these sequences formed an independent, disorganized clade (Figure 5A clade containing yellow and blue sequences), relative to those recovered from dinoflagellate genomes or other invertebrate hosts. *MCP* and *RdRp* ORFs putatively derived from the same dinoflagellate genomes often shared clades (clades containing multiple blue or green sequences in Figure 5A, B), perhaps indicative of duplications within genomes or multiple integration events of particular dinoRNAV lineages within host genera. Symbiodiniaceae first diversified from the psychrophilic, free-living *Polarella* outgroup ~160 million years ago (Janouškovec et al. 2017, Stephens et al. 2020; LaJeunesse et al. 2018). The detection of putative dinoRNAV *RdRp* ORFs within *Polarella* genomes is therefore indicative of either the antiquity of dinoRNAV-dinoflagellate interactions and/or a propensity for dinoRNAV integration across Dinophyceae families. However, the exclusion of the *P. glacialis* dinoRNAV-*RdRp* from *RdRps* of other dinoflagellate clades (pink, Figure 5B) further illustrates the congruence between EVEs and their host genomes.

**Figure 5.**
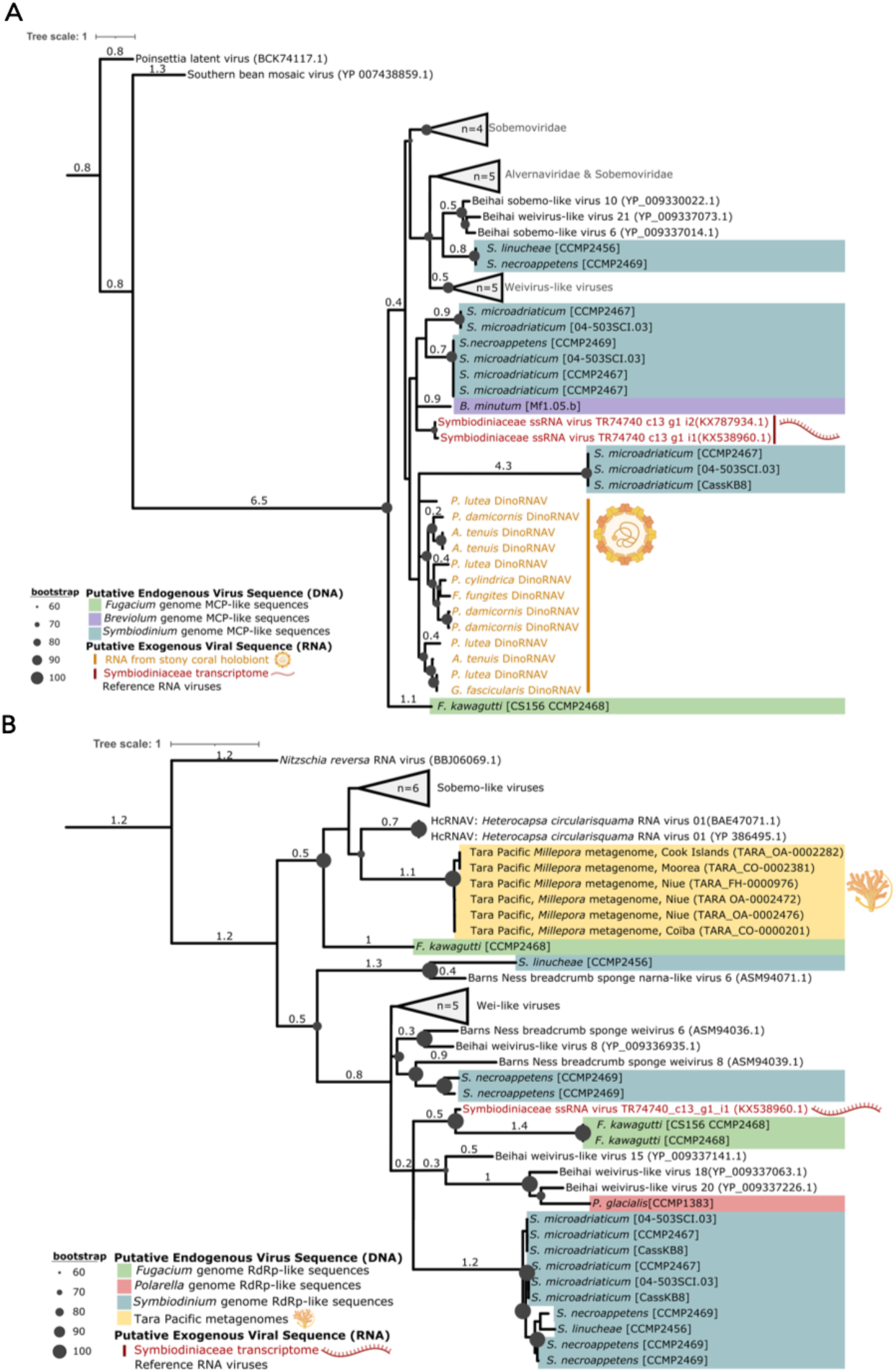
Phylogenies of dinoRNAV Major Capsid Protein (*MCP*, **A**) and RNA-dependent RNA polymerase (*RdRp*, **B**) ORFs recovered from host metagenomes, transcriptomes, polished genomes, and extant +ssRNA reference viruses from amplicon libraries (A only). Both trees include *Dinornavirus* reference sequences. (A) Maximum-likelihood tree of *MCP* amino acid sequences generated with a LG+F+G4 substitution model and 50,000 parametric bootstraps, illustrating the similarity of putative dinoRNAV EVEs (this study) to extant dinoRNAVs from stony coral colonies. (B) Maximum-likelihood tree of *RdRp* amino acid sequences generated with a Blosum62+G4 substitution model and 50,000 parametric bootstraps, demonstrating the similarity of metagenomic dinoRNAV EVE *RdRps* to *RdRps* of the sole recognized *Dinornavirus*, Heterocapsa circularisquama RNA virus (HcRNAV), as well as alignment to each other. Trees were visualized in iTOL. *[this image is a low resolution placeholder]*

The expression and functional potential of endogenized dinoRNAVs (if any) remains unclear. Two transcripts derived from *Cladocopium* transcriptomes and annotated as *MCP* ORFs of +ssRNA viral sequences (‘TR74740_c13-g1_i1’ and ‘TR74740_c13-g1_i2’, Levin et al. 2017, red text in Figure 5A) shared a clade with putative *Symbiodinium* dinoRNAV EVEs. Likewise, the *RdRp* ORF of ‘TR74740_c13-g1_i1’ and the *RdRp* of ‘GAKY01194223.1’— a transcript derived from a cultured *Symbiodinium microadriaticum* A1 transcriptome—shared similarity to putative endogenous dinoRNAVs (Figure 5B; Levin et al. 2017, Baumgarten et al. 2013). Importantly, both RNA transcripts also shared features characteristic of dinoflagellates, such as a 5′ spliced leader sequence (“DinoSL”; Zhang et al. 2013) or dinoflagellate sequence space flanking the dinoRNAV itself (Baumgarten et al. 2013). While ‘TR74740_c13-g1_i1’ appeared to be in the top 0.03% of expressed transcripts at under certain thermal conditions, GAKY01194223.1 appeared exhibit moderately differential expression at the extremes of temperature and ionic stress in a cultured host (Levin et al. 2017, Baumgarten et al. 2013).

While viral *RdRps* have been leveraged by eukaryotes in multiple pathways (Lipardi and Paterson 2010), the fragmented nature of the putative dinoRNAV EVEs *in silico* may contribute to a role in triggering antiviral mechanisms in their hosts (Blair et al. 2020, Suzuki et al. 2020). Given that the *Symbiodinium* genome contains all core RNAi protein machinery, including Argonaute and Dicer, and that GAKY01194223.1 folds into several hairpins (ΔG = - 142.5kcal/mol; Supplemental Figure 4 examples), Symbiodiniaceae may use the putative EVE ncRNA identified here to develop host immunity against extant, exogenous dinoRNAVs.

Furthermore, *S. microadriaticum* harboring dinoRNAV EVEs contained numerous non-retroviral EVEs of other viral families (Supplemental Figure 5) in close proximity, such as *Herpesviridae, Baculoviridae, Poxviridae, Iridoviridae, Phycodnaviridae, Pandoraviridae* and *Pithoviridae*, ssDNA viruses of the family *Shotokuvirae*, -ssRNA viruses from the family *Rhabdoviridae* and +ssRNA viruses from the family *Coronaviridae* (Supplemental Figure 5). Tara Pacific metagenomes corroborate findings of similar *RdRps* from these viral families (Supplemental Figure 5).

## Conclusions

Endogenized viral elements, such as dinoRNAVs, demonstrate how *in silico* identification can provide context for viral genomes in non-model, symbiotic systems such as coral holobionts, impacting how we study coral reefs and their viral consortia. Endogenized viral elements (EVEs) have been effectively utilized in many terrestrial systems to better understand the evolutionary history of viruses (“paleovirology”) and pair hosts with extant viruses in multipartite systems. We propose that dinoRNAVs utilize host-provisioned mechanisms (e.g., LINEs) to integrate into single-celled dinoflagellate genomes as EVEs. We detected heritable integrations of multiple putative dinoRNAV genes in Symbiodiniaceae scaffolds from cnidarian metagenomes, as well as in diverse genomes of cultured Symbiodiniaceae; no integrations were detected from seawater metagenomes nor diverse aposymbiotic cnidarian genomes. The apparent pervasive nature of dinoRNAV-like sequences among dinoflagellate genomes (especially the genus *Symbiodinium*) suggests widespread and recurrent/ancestral integration and conservation of these EVEs. The findings presented in this study further validate the dinoRNAV-Symbiodiniaceae virus-host pair, enhancing our understanding of ecologically and economically important cnidarian holobionts and opening the door to examining the role of EVEs in reef health.

## Data Availability

Metadata and sequences are accessible in zenodo: https://zenodo.org/communities/tarapacific?page=1&size=20

## Acknowledgements and Contributions

Special thanks to the Tara Ocean Foundation, the R/V Tara crew and the Tara Pacific Expedition Participants (https://doi.org/10.5281/zenodo.3777760). We are keen to thank the commitment of the following institutions for their financial and scientific support that made this unique Tara Pacific Expedition possible: CNRS, PSL, CSM, EPHE, Genoscope, CEA, Inserm, Université Côte d’Azur, ANR, agnès b., UNESCO-IOC, the Veolia Foundation, the Prince Albert II de Monaco Foundation, Région Bretagne, Billerudkorsnas, AmerisourceBergen Company, Lorient Agglomération, Oceans by Disney, L’Oréal, Biotherm, France Collectivités, Fonds Français pour l’Environnement Mondial (FFEM), Etienne Bourgois, and the Tara Ocean Foundation teams. Tara Pacific would not exist without the continuous support of the participating institutes. This research is further supported by NSF OCE #1635798 to AMSC, NSF DOB Grant 2025457 to RLVT, and with additional support from NSF PRFB – 1907184 to KSIB.

## List of Tara Pacific Consortium Coordinators

Sylvain Agostini (orcid.org/0000-0001-9040-9296) Shimoda Marine Research Center, University of Tsukuba, 5-10-1, Shimoda, Shizuoka, Japan

Denis Allemand (orcid.org/0000-0002-3089-4290) Centre Scientifique de Monaco, 8 Quai Antoine Ier, MC-98000, Principality of Monaco

Bernard Banaigs (orcid.org/0000-0003-3473-4283) PSL Research University: EPHE-UPVD- CNRS, USR 3278 CRIOBE, Université de Perpignan, France

Emilie Boissin (orcid.org/0000-0002-4110-790X) PSL Research University: EPHE-UPVD- CNRS, USR 3278 CRIOBE, Laboratoire d’Excellence CORAIL, Université de Perpignan, 52 Avenue Paul Alduy, 66860 Perpignan Cedex, France

Emmanuel Boss (orcid.org/0000-0002-8334-9595) School of Marine Sciences, University of Maine, Orono, 04469, Maine, USA

Chris Bowler (orcid.org/0000-0003-3835-6187) Institut de Biologie de l’Ecole Normale Supérieure (IBENS), Ecole normale supérieure, CNRS, INSERM, Université PSL, 75005 Paris, France

Colomban de Vargas (orcid.org/0000-0002-6476-6019) Sorbonne Université, CNRS, Station Biologique de Roscoff, AD2M, UMR 7144, ECOMAP 29680 Roscoff, France & Research Federation for the study of Global Ocean Systems Ecology and Evolution, FR2022/ Tara Oceans-GOSEE, 3 rue Michel-Ange, 75016 Paris, France

Eric Douville (orcid.org/0000-0002-6673-1768) Laboratoire des Sciences du Climat et de l’Environnement, LSCE/IPSL, CEA-CNRS-UVSQ, Université Paris-Saclay, F-91191 Gif-sur-Yvette, France

Michel Flores (orcid.org/0000-0003-3609-286X) Weizmann Institute of Science, Department of Earth and Planetary Sciences, 76100 Rehovot, Israel

Didier Forcioli (orcid.org/0000-0002-5505-0932) Université Côte d’Azur, CNRS, INSERM, IRCAN, Medical School, Nice, France and Department of Medical Genetics, CHU of Nice, France

Paola Furla (orcid.org/0000-0001-9899-942X) Université Côte d’Azur, CNRS, INSERM, IRCAN, Medical School, Nice, France and Department of Medical Genetics, CHU of Nice, France

Pierre Galand (orcid.org/0000-0002-2238-3247) Sorbonne Université, CNRS, Laboratoire d’Ecogéochimie des Environnements Benthiques (LECOB), Observatoire Océanologique de Banyuls, 66650 Banyuls sur mer, France

Eric Gilson (orcid.org/0000-0001-5738-6723) Université Côte d’Azur, CNRS, Inserm, IRCAN, France

Fabien Lombard (orcid.org/0000-0002-8626-8782) Sorbonne Université, Institut de la Mer de Villefranche sur mer, Laboratoire d’Océanographie de Villefranche, F-06230 Villefranche-sur-Mer, France

Stéphane Pesant (orcid.org/0000-0002-4936-5209) European Molecular Biology Laboratory, European Bioinformatics Institute, Wellcome Genome Campus, Hinxton, Cambridge CB10 1SD, UK

Serge Planes (orcid.org/0000-0002-5689-5371) PSL Research University: EPHE-UPVD-CNRS, USR 3278 CRIOBE, Laboratoire d’Excellence CORAIL, Université de Perpignan, 52 Avenue Paul Alduy, 66860 Perpignan Cedex, France

Stéphanie Reynaud (orcid.org/0000-0001-9975-6075) Centre Scientifique de Monaco, 8 Quai Antoine Ier, MC-98000, Principality of Monaco

Matthew B. Sullivan (orcid.org/0000-0003-4040-9831) The Ohio State University, Departments of Microbiology and Civil, Environmental and Geodetic Engineering, Columbus, Ohio, 43210 USA

Shinichi Sunagawa (orcid.org/0000-0003-3065-0314) Department of Biology, Institute of Microbiology and Swiss Institute of Bioinformatics, Vladimir-Prelog-Weg 4, ETH Zürich, CH-8093 Zürich, Switzerland

Olivier Thomas (orcid.org/0000-0002-5708-1409) Marine Biodiscovery Laboratory, School of Chemistry and Ryan Institute, National University of Ireland, Galway, Ireland

Romain Troublé (ORCID not-available) Fondation Tara Océan, Base Tara, 8 rue de Prague, 75 012 Paris, France

Rebecca Vega Thurber (orcid.org/0000-0003-3516-2061) Oregon State University, Department of Microbiology, 220 Nash Hall, 97331Corvallis OR USA

Christian R. Voolstra (orcid.org/0000-0003-4555-3795) Department of Biology, University of Konstanz, 78457 Konstanz, Germany

Patrick Wincker (orcid.org/0000-0001-7562-3454) Génomique Métabolique, Genoscope, Institut François Jacob, CEA, CNRS, Univ Evry, Université Paris-Saclay, 91057 Evry, France

Didier Zoccola (orcid.org/0000-0002-1524-8098) Centre Scientifique de Monaco, 8 Quai Antoine Ier, MC-98000, Principality of Monaco

